# scGPD: single-cell informed gene panel design for targeted spatial transcriptomics

**DOI:** 10.1101/2025.10.02.680117

**Authors:** Yunshan Guo, Jia Zhao, Rui B. Chang, Hongyu Zhao

## Abstract

In targeted spatial transcriptomics technologies, a key challenge is to select an informative gene panel that captures the complexity of cellular and spatial heterogeneity within tissues. Many existing methods use prior knowledge or heuristic selection rules, such as selecting highly variable genes, which overlook gene-gene correlations and may consequently result in suboptimal coverage. To address the limitations of the existing methods, we introduce scGPD, a deep learning-based framework for gene panel design that leverages single-cell RNA-seq data to identify compact, nonredundant sets of genes for spatial profiling. scGPD uses a gene-gene correlation-aware gating mechanism to extract informative features from data, encouraging diversity among selected genes and eliminating redundancy. Across diverse single-cell datasets, scGPD outperforms existing gene panel design methods in recovering transcriptome-wide expression using a limited number of genes. When applied to spatial transcriptomics data, it achieves superior cell type classification accuracy, demonstrating strong generalization across modalities. The gene panels selected by scGPD further exhibit well-defined spatial expression patterns, highlighting their robustness and relevance for spatial analysis. The scGPD framework is flexible and can be adapted to multiple use cases, enabling the prioritization of genes relevant to specific diseases or phenotypes. Together, these results demonstrate that scGPD provides a robust and adaptable solution to design efficient gene panels for spatial transcriptomics, with broad applicability to tissue mapping and disease characterization.

## Introduction

Single-cell RNA sequencing (scRNA-seq) has revolutionized the study of cellular heterogeneity by enabling transcriptomic profiling at single-cell resolution across diverse tissues and biological systems [1] [2] [3] [4]. This technology has been instrumental in uncovering novel cell types, developmental trajectories, and disease-associated transcriptional programs. However, a fundamental limitation of scRNA-seq is the loss of spatial context, as tissues must be dissociated into single-cell suspensions. This dissociation disrupts native cell–cell interactions and obliterates spatial patterns of gene expression that are essential for understanding tissue organization and microenvironmental influences [5].

To address this shortcoming, spatial transcriptomics (ST) technologies have been developed to enable spatially resolved gene expression profiling within intact tissue sections. These methods preserve tissue architecture while allowing for the detection of mRNA abundance at various spatial resolutions [6] [7] [8] [9]. ST technologies can be broadly categorized into two categories: spot-based ST (spot-ST) and single-cell ST (sc-ST) approaches. Spot-ST methods, such as 10X Visium [10] and HDST[11], quantify gene expression within spatially resolved spots. While these approaches provide transcriptome-wide coverage, they lack single-cell resolution, making them less suitable for cell-type specific analyses, such as studying spatial variation in ligand–receptor interactions. In contrast, sc-ST technologies, such as MERFISH [9], seqFISH, seqFISH+ [12] and osmFISH[8], provide subcellular spatial precision and single-molecule sensitivity, which makes them particularly well suited for high-resolution tissue mapping [5]. Despite these advantages, being fluorescence in situ hybridization (FISH)-based, each sc-ST dataset is limited by the number of genes that can be simultaneously measured, typically in the range of a few hundred to about a thousand genes, which restricts the full potential of sc-ST data in capturing comprehensive transcriptomic landscapes. Consequently, the design of targeted gene panels is a critical step in sc-ST workflows. The selection of a biologically informative and functionally relevant subset of genes is essential to accurately capture the cellular diversity and spatial organization within tissues.

Gene selection for targeted spatial transcriptomics is frequently guided by heuristic strategies, such as choosing well-characterized marker genes or those with high expression in specific cell subsets. Although these approaches are straightforward, they often fail to capture genes with more subtle, heterogeneous, or spatially complex expression patterns. A critical limitation of such methods is their tendency to overlook correlations between genes, an issue that can significantly compromise the informativeness of the selected panel. When co-expressed or functionally related genes are selected together, the resulting panel may include redundant signals that do not contribute additional discriminatory power. This redundancy not only reduces the overall efficiency of the panel, but also limits its ability to capture the full diversity of cell types or spatial domains present in the tissue [13] [14] [15].

To address these challenges, we reformulate gene selection as a feature selection problem and propose scGPD (single-cell informed Gene Panel Design), a principled, data-driven framework grounded in deep learning. The framework leverages single-cell RNA-seq data to guide panel selection, systematically prioritizing genes that are informative and nonredundant to yield compact panels that preserve biological signal.

A key innovation of scGPD lies in its correlation-aware gating mechanism [16], which explicitly accounts for gene–gene dependencies during training. Rather than allowing highly correlated genes to be selected redundantly, our gating strategy promotes competition among correlated features, effectively driving the model to prioritize the most informative representative within each correlated cluster. This results in gene panels being optimized for downstream tasks such as transcriptome reconstruction or cell type classification. In contrast to existing methods, scGPD is also adaptable: it can scale to large datasets via minibatched training and accommodate diverse experimental designs and panel size requirements.

We demonstrate the efficacy and versatility of scGPD through extensive benchmarking on multiple publicly available single-cell RNA-seq datasets, covering a wide range of dataset sizes and cellular diversity. Across these datasets, scGPD consistently selects informative genes that preserve key biological variation. Beyond quantitative benchmarking, we show that the genes selected by scGPD from single-cell data also exhibit coherent spatial expression patterns in the corresponding spatial transcriptomics datasets, highlighting the method’s ability to identify biologically meaningful markers of spatial relevance. Furthermore, we demonstrate that scGPD can be adapted to prioritize disease-associated genes by modifying the objective of feature selection, enabling the targeted discovery of candidate biomarkers for spatial profiling in pathological contexts.

## Results

### Method Overview

scGPD is a deep learning framework that leverages reference single-cell RNA-seq data to construct gene panels optimized for specific experimental objectives. Through its specialized architecture with differentiable feature selection layers, the method identifies a compact set of highly informative genes that effectively capture transcriptome-wide expression patterns and preserve biologically meaningful cellular variation.

The framework consists of two sequential stages (Figure 1). In the first stage, scGPD reconstructs transcriptome-wide expression profiles from a reduced set of candidate genes. Rather than evaluating genes independently, the model employs a correlation-aware binary gating mechanism that identifies interdependent gene groups using a Gaussian copula to model gene dependencies [17]. This gating mechanism utilizes the Binary Concrete distribution [18] to select the most informative representative from each correlated group while eliminating redundant genes, resulting in a narrowed gene pool containing the most salient candidate features. To appropriately match the distributional characteristics of scRNA-seq data, scGPD employs Poisson loss as its objective function [19, 20, 21, 22], which handles the count-based nature of single-cell expression data. In the second stage, scGPD further refines the gene selection by choosing exactly *k* genes from the reduced candidate set (of size *d*) obtained from the first stage. This refinement again employs a binary gating mechanism, but now uses masks derived from Concrete distributions [18] to enforce a strict selection budget of exactly *k* genes. To ensure that the final gene panel is optimized for specific biological applications, the model employs task-specific loss functions tailored to downstream objectives, such as cell type classification or spatial expression prediction.

**Figure 1.**
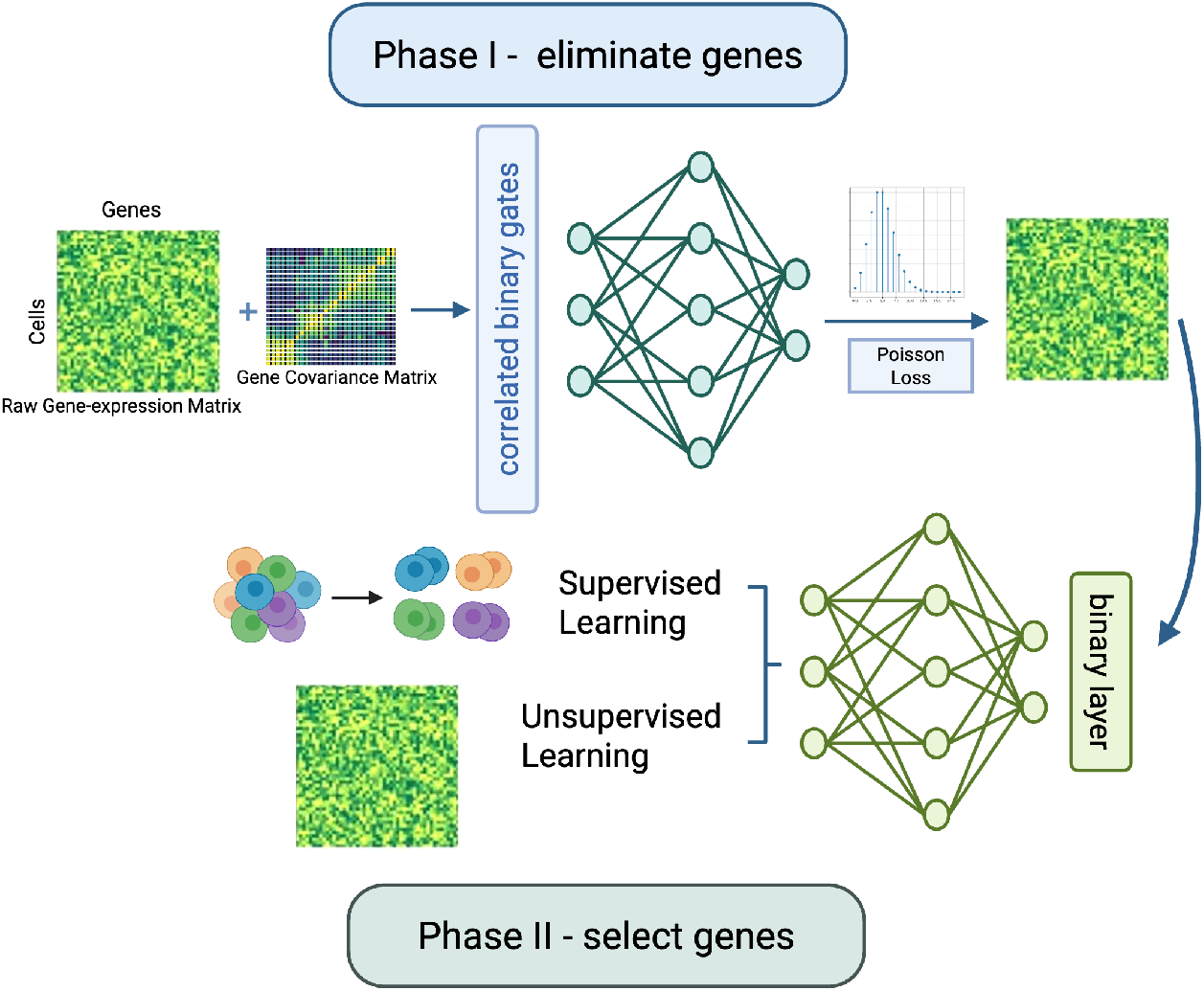
Overview of scGPD. The scGPD framework implements a dual-stage feature selection paradigm for identifying informative genetic markers. In the first stage, to reconstruct the original scRNA-seq gene expression levels, a correlated binary gating mechanism is employed to eliminate redundant and uninformative genes by learning interdependent feature activations. In the second stage, application-specific loss functions guide the selection of exactly k genes from the reduced pool of d candidates, achieved by applying a binary mask to the model inputs.

### scGPD enables accurate cell type annotation and scRNA-seq expression profiles reconstruction

We evaluated scGPD on three scRNA-seq datasets derived from distinct biological sources: pancreas [23], heart [24], and peripheral blood mononuclear cells (PBMC) [4]. Starting from an initial set of 5,000 highly variable genes and the gene covariance estimated from CS-CORE[25], we applied scGPD and three baseline gene selection methods: PERSIST [26], scGIST [27], and geneBasis [28] to identify gene panels ranging in size from 32 to 256 genes, a range that encompasses the majority of panel sizes used in FISH experiments.

As a flexible analytical framework, scGPD supports both supervised and unsupervised learning paradigms, making it applicable to a wide range of downstream tasks. To evaluate its effectiveness, we first assessed how well selected gene panels could reconstruct the original scRNA-seq expression profiles. Specifically, we computed Pearson correlation coefficients [29] between the original and reconstructed gene expression matrices across genes for each cell. To ensure a fair comparison, we trained the same multilayer perceptron (MLP) network architecture to map each method’s selected gene panels to the full expression profiles. Across all datasets and panel sizes, scGPD consistently achieved higher reconstruction accuracy than competing methods (Figure 2A). These results underscore the robustness and reliability of scGPD for guiding targeted sequencing.

**Figure 2.**
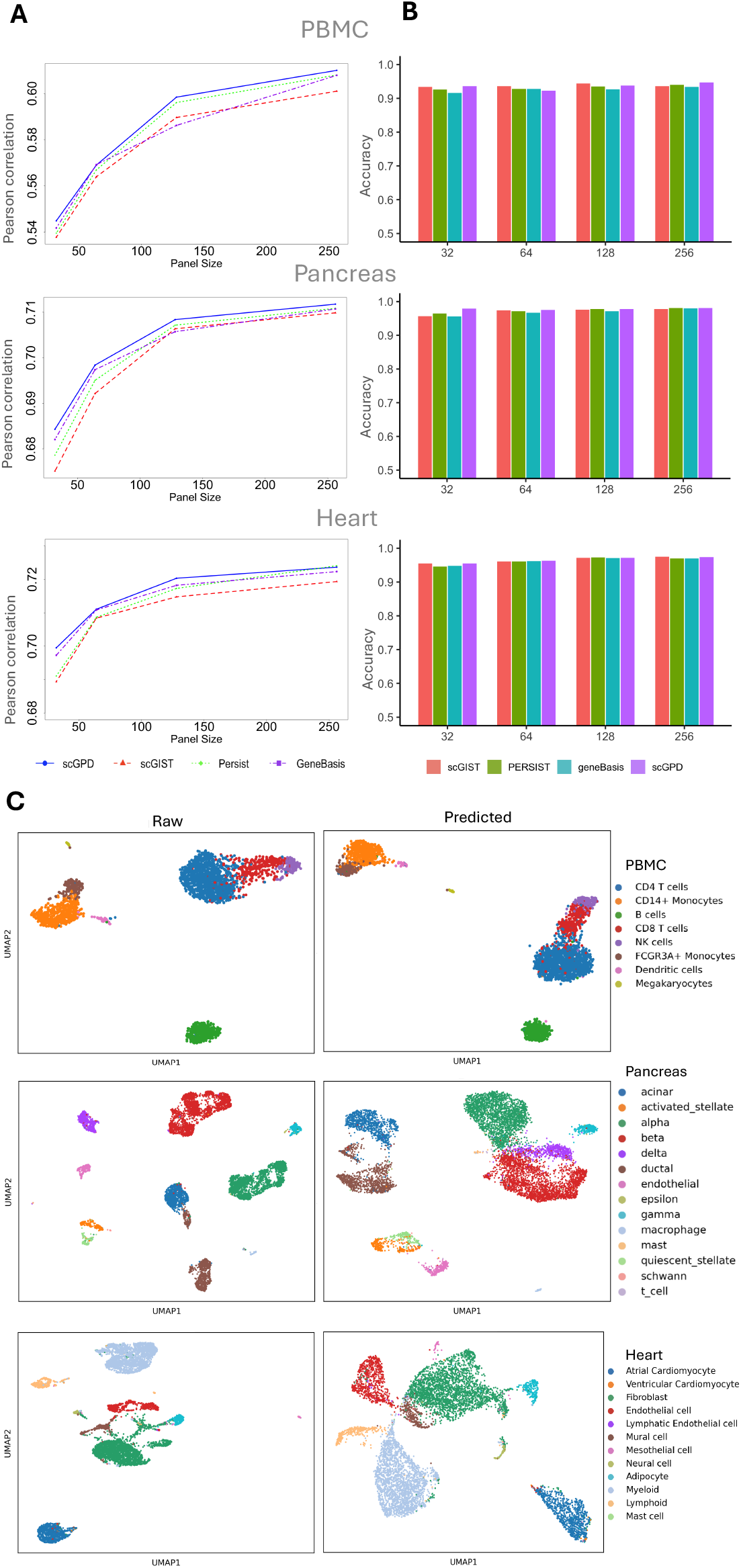
Performance of various methods on three different datasets. **A**. Pearson correlation coefficients for variable panel sizes. **B**. Accuracies for variable panel sizes. **C**. UMAP plots using the gene panel of size 256 selected by scGPD for the t1h8ree datasets.

We further evaluated the utility of selected gene panels in a cell-type classification task, a critical downstream analysis in many single-cell transcriptomics studies. For each method, we trained the same MLP classifiers using the gene panels selected by the respective methods and assessed classification accuracy across varying panel sizes. As expected, larger gene panels generally led to improved classification performance across all methods, reflecting the increased information content available for discriminating cell types. Overall, scGPD achieved classification performance that was comparable to or better than existing state-of-the-art methods across most conditions and it consistently maintained competitive accuracy across different gene panel sizes (Figure 2B, Supplementary Figure S1). This demonstrates that scGPD consistently identifies informative gene subsets that perform well in supervised learning contexts across different gene panel sizes. We visualizes the UMAP embedding of the three datasets derived based on raw data and the 256-gene panel selected by scGPD. The resulting projection reveals that cells cluster according to their annotated cell types, demonstrating strong concordance between the selected gene panel and the underlying transcriptional structure (Figure 2C). This alignment suggests that scGPD effectively captures cell-type discriminative features, enabling accurate low-dimensional representations that preserve biological identity. These results underscore the versatility of scGPD and its robustness across different analytical scenarios, demonstrating its potential as a reliable tool for gene panel design in resource-limited settings.

Collectively, these results demonstrate that scGPD is a powerful and versatile method for gene panel selection, achieving competitive or superior performance on multiple evaluation metrics while remaining flexible across various applications.

### Performance evaluation on a spatial transcriptomics dataset

scGPD can identify informative marker genes for a variety of experimental objectives. Here, we applied the genes selected using scRNA-seq to data collected from spatial transcriptomics studies. We analyzed a seqFISH+ dataset [12] from mouse olfactory, where a corresponding scRNA-seq dataset [30] is also available. To ensure consistency, we restricted our analysis to the 9,913 common genes present in both datasets. We then used scGPD to generate gene panels from the scRNA-seq data and predicted cell types in the seqFISH+ dataset based on these panels. The seqFISH+ dataset’s pre-existing cell-type annotations served as ground truth labels, enabling quantitative assessment of how effectively scGPD-selected gene panels preserve cell-type discrimination when transferred from single-cell to spatial transcriptomics contexts.

We benchmark the performance of scGPD with the other three methods: scGIST, geneBasis and PERSIST. Although scGPD shows slightly lower performance for smaller panels, when panel size increases, scGPD outperforms the other methods, achieving the highest accuracy at larger panel sizes (Figure 3A). PERSIST and scGIST exhibit similar trends and GeneBasis is the most competitive baseline method. Figure 3B presents the confusion matrix of predicted cell by scGPD using a 256-gene panel, evaluated on the seqFISH+ test dataset.

**Figure 3.**
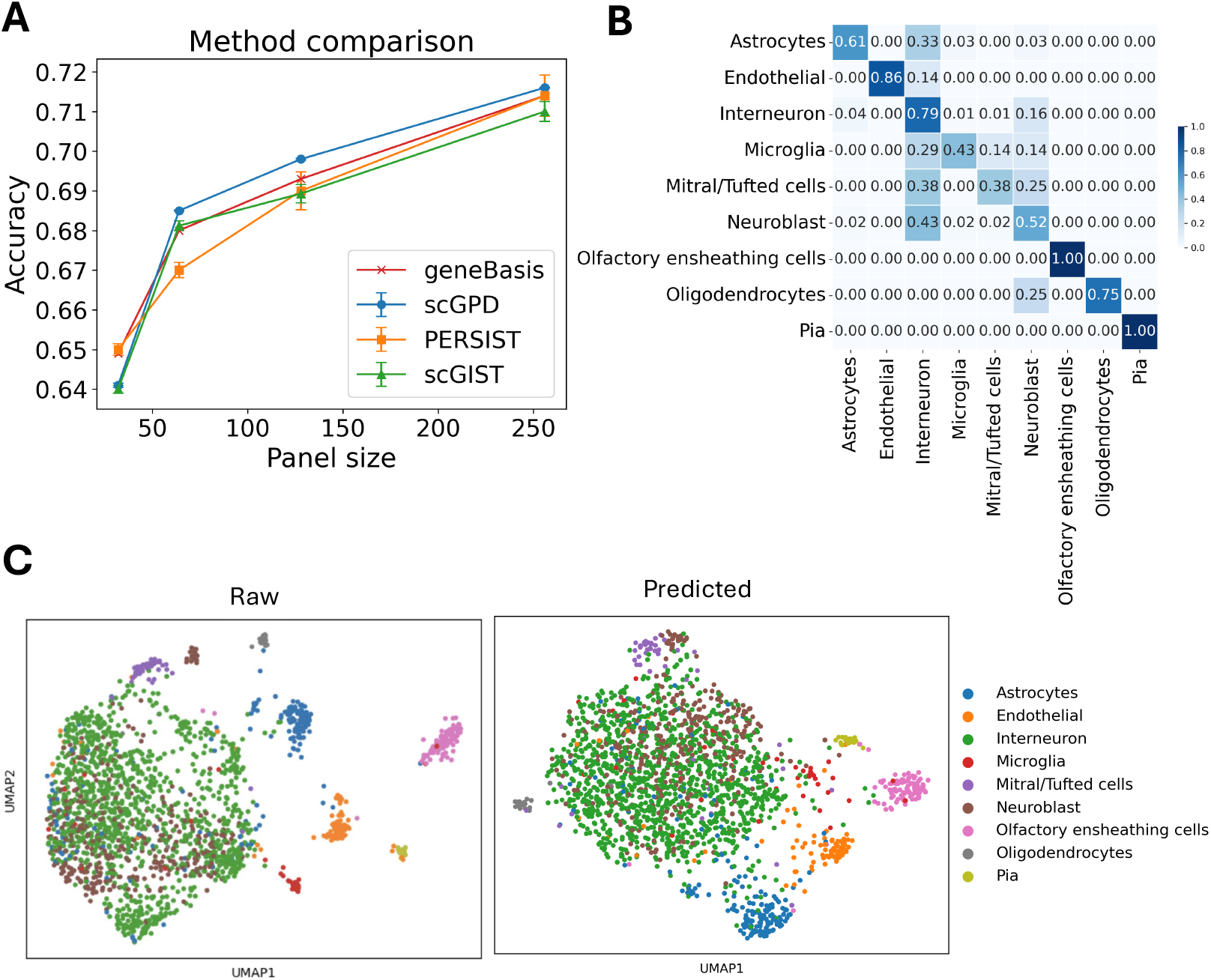
Performance evaluation of gene panel selection on a spatial transcriptomics dataset. **A**. Cell type classification accuracy using gene panels designed from scRNA-seq data and evaluated on the seqFISH+ test set. **B**. Confusion matrix of cell type predictions by scGPD using a panel of 256 genes, evaluated on the seqFISH+ test dataset. **C**. UMAP visualization of the seqFISH+ data derived based on raw data and the 256-gene panel selected by scGPD, demonstrating alignment between predicted cell types and the original clustering structure.

Supplementary Figure S2 presents the PAGA similarity [31] scores for varying-size gene panels (32 to 256 genes), evaluating the ability of each method to preserve the structure of cell-cell similarity. Across all panel sizes, scGPD demonstrates consistently strong performance, maintaining high PAGA similarity scores and outperforming scGIST and PERSIST at all panel sizes. This pattern underscores the ability of scGPD to select gene panels that faithfully retain the structure of the cellular manifold. Using the 256 genes identified from scGPD, we show a UMAP projection derived based on raw data and selected gene set to compare the cell-type distributions (Figure 3C). The predicted UMAP maintains the overall structure of the raw dataset, demonstrating that our model successfully captures the differences among cell types. Together, these results illustrate the effectiveness of scGPD in selecting informative gene panels that generalize well to spatial transcriptomics data.

### Identifying informative genes reveals spatially distinct expression patterns in spatial transcriptomics data

Our proposed method, scGPD, effectively identifies genes with spatially distinct expression patterns in spatial transcriptomics data. Using a human breast cancer 10x Genomics scRNA-seq dataset [32] as a reference, we applied scGPD-identified genes to data from breast cancer tissue [32] profiled by 10x Visium to explore their role in dissecting the tumor microenvironment. The selected genes exhibit clear spatial variation, with specific regions showing elevated expression, indicating their significance in defining distinct tissue microenvironments and potentially influencing tumor heterogeneity (Figure 4A).

**Figure 4.**
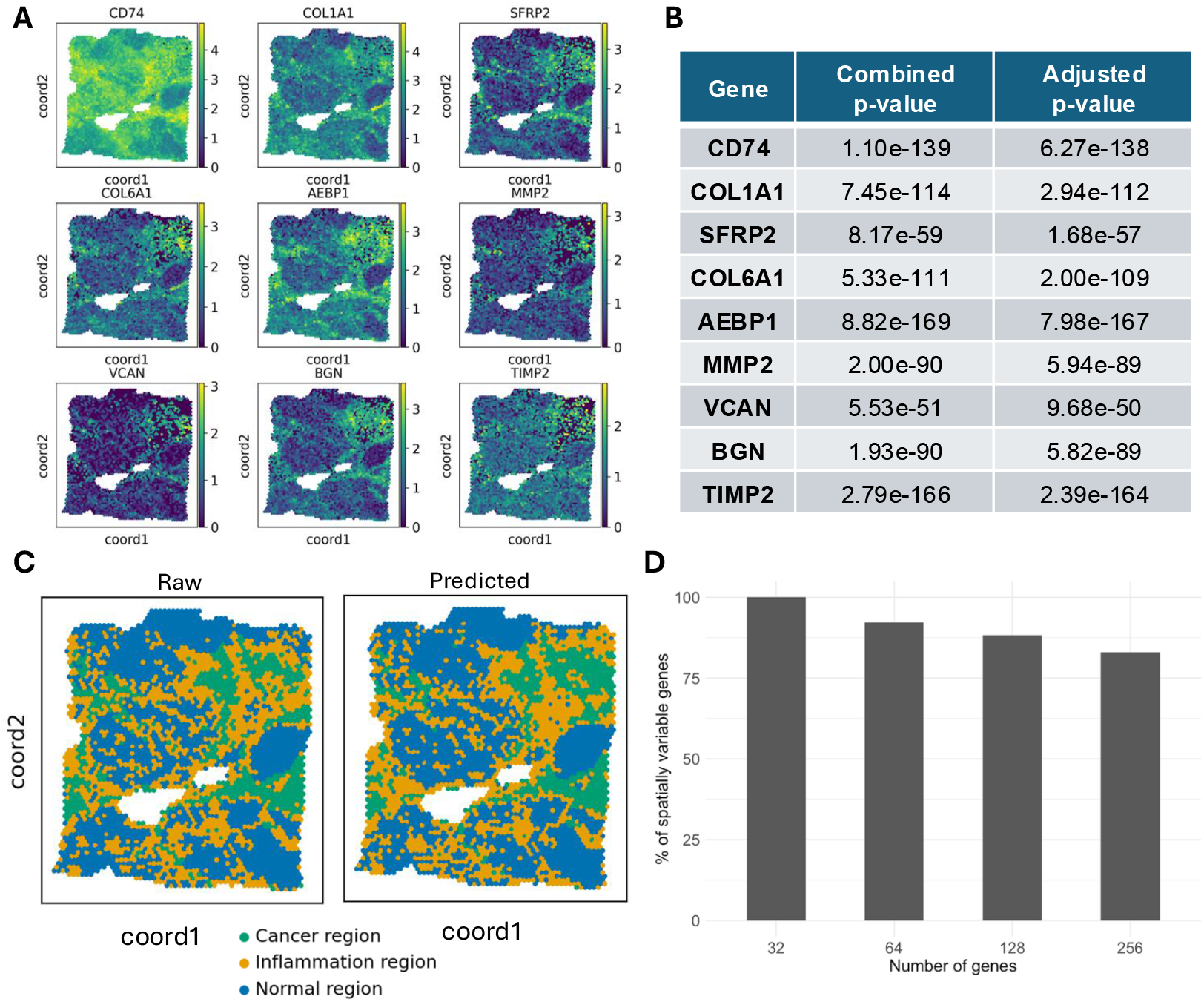
Spatial validation and functional relevance of scGPD-selected genes in breast cancer tissue. **A**. Spatial expression patterns of representative scGPD-identified genes in 10x Visium breast cancer ST data, showing clear spatially distinct expression domains. **B**. Statistical validation of spatial variability using SPARK, with significant adjusted p-values confirming spatial expression of selected genes. **C**. Tissue segmentation using scGPD-selected genes compared to histological annotations from transcriptome-wide profiles, demonstrating strong concordance and preservation of tumor architecture. **D**. Bar plot showing the proportion of spatially variable genes retained across increasing gene panel sizes.

To statistically evaluate the spatial variability of these genes, we employed SPARK [19], a robust statistical framework for detecting spatially expressed genes. The analysis (Figure 4B, Supplementary Table S1-S4) revealed highly significant combined and adjusted p-values, confirming the meaningful spatial expression of the identified genes and reinforcing the robustness of scGPD in capturing genes relevant to spatial tissue organization.

Importantly, the genes identified, such as CD74, COL1A1, SFRP2, and COL6A1, have wellestablished biological relevance in breast cancer. CD74 promotes triple-negative breast cancer progression by expanding immunosuppressive cells [33]. COL6A1 facilitates breast cancer growth and spread by supporting tumor cell proliferation and creating a favorable tumor environment and COL1A1 is a structural protein in the extracellular matrix and is involved in cancer spread and metastasis [34, 35]. The spatial expression patterns of these genes likely reflect critical biological processes in the tumor microenvironment, such as immune cell localization, stromal activation, and epithelial-mesenchymal transitions. Together, these findings underscore the ability of scGPD to identify genes that are not only statistically significant in their spatial distribution but also biologically meaningful in the context of cancer progression and tissue heterogeneity. We also evaluated the impact of gene panel size on spatial variability detection. By systematically varying panel sizes from 32 to 256, we assessed how the proportion of spatially variable genes changes. The results indicate that even with an expanded gene set, scGPD retains at least 80% of spatially variable genes (adjusted p-value *<* 0.05), demonstrating its ability to efficiently prioritize informative genes while maintaining biological interpretability.

To further assess the functional utility of the identified genes, we used the 256 genes for tissue segmentation and compared the results to ground truth histological annotations. These ground truth annotations were established by first predicting the spatial distribution of cell types in the ST data through integration with corresponding scRNA-seq data, then dividing the tissue section into three distinct tissue regions based on the spatial distribution patterns of these cell types. The strong concordance between the predicted and reference segmentations (Figure 4C), with an adjusted Rand index (ARI) of 0.63, highlights the ability of scGPD to accurately capture spatially distinct biological regions.

These findings underscore the power of scGPD in optimizing gene selection for spatial transcriptomics, enhancing tissue characterization, and enabling deeper insights into spatial gene expression. By identifying genes with strong spatial signals, scGPD provides a scalable and efficient solution for uncovering the spatial organization of complex tissues.

### Discovery of disease-related genes for lung adenocarcinoma (LUAD)

scGPD is a powerful and versatile tool for uncovering disease-related genes by systematically analyzing scRNA-seq data from samples under different conditions. By leveraging its ability to distinguish gene expression variations at the single-cell level, scGPD enables the identification of key molecular signatures associated with disease progression. This capability is particularly valuable for studying complex diseases where cellular heterogeneity plays a crucial role in pathogenesis.

In this study, we apply scGPD to LUAD, a highly prevalent and aggressive subtype of non-small cell lung cancer (NSCLC). LUAD is characterized by tumor heterogeneity and clinical symptoms such as persistent coughing, chest pain, and respiratory decline. Understanding its underlying genetic risk factors is essential for unraveling its molecular mechanisms and developing targeted therapeutic strategies.

To comprehensively investigate LUAD-specific gene expression patterns, we analyzed a single-cell RNA-seq dataset [36] comprising LUAD and healthy control (HC) samples. Figure 5A presents a heatmap of gene expression across normal and tumor samples, revealing key differentially expressed genes (DEGs) that may serve as LUAD biomarkers [37] [38] [39]. Using scGPD in a supervised setting, we identified disease-related genes by training the model to distinguish between HC and LUAD samples based on single-cell transcriptomic profiles. The genes selected by scGPD were those most informative for accurately classifying cells by disease status. For example, among the highlighted genes, FABP4 modulates lipid metabolism and immune responses by influencing NK and macrophage function [40]; MMP7 facilitates tumor proliferation and metastasis through matrix degradation and angiogenesis [41]; and C1QB contributes to immune microenvironment regulation via the complement cascade [42]. These findings provide insights into tumor microenvironment heterogeneity and highlight potential driver genes contributing to LUAD pathogenesis.

**Figure 5.**
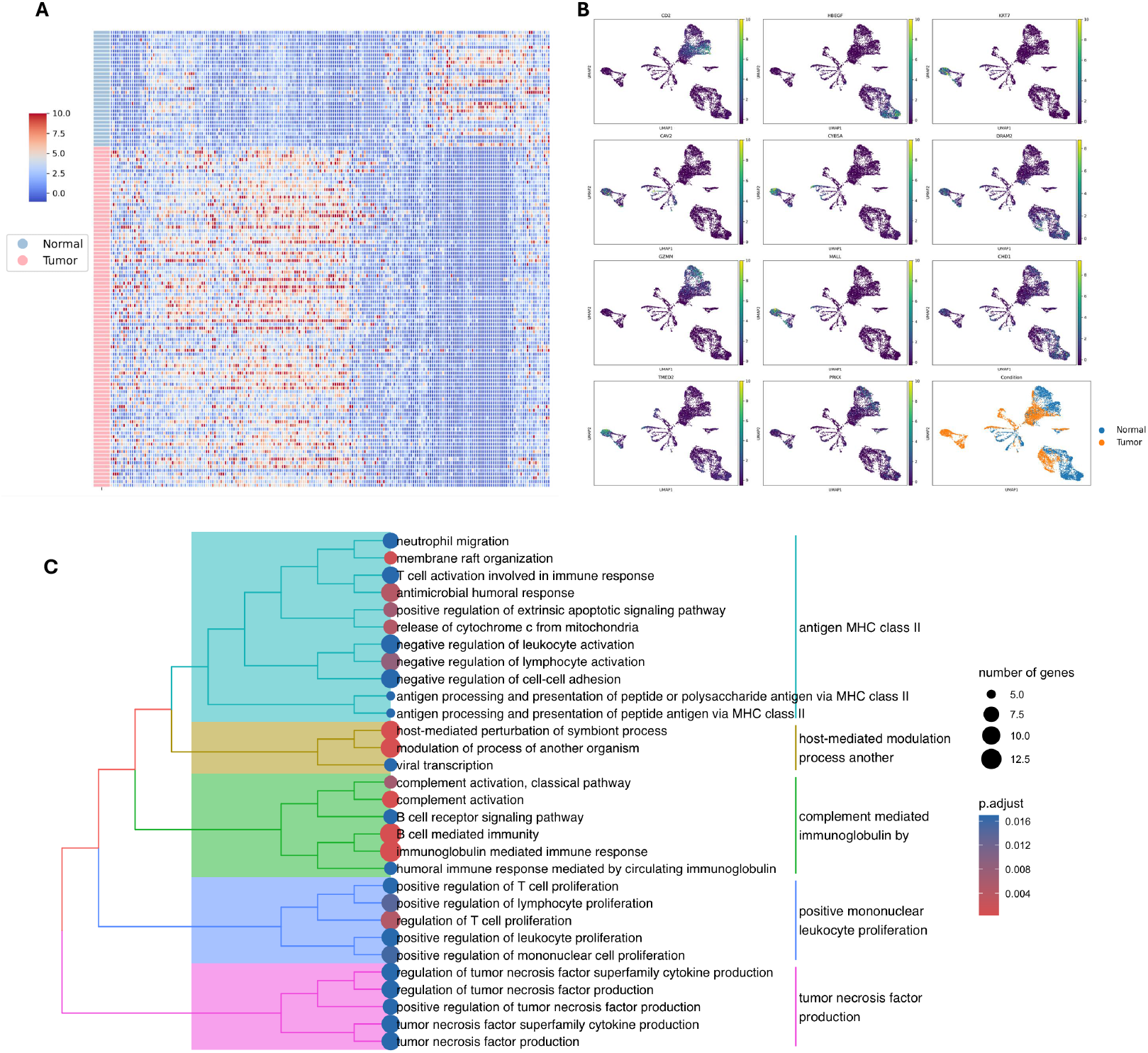
Identification of LUAD-related genes and their expression dynamics across cell types. **A**. Heatmap showing differentially expressed genes between healthy control (HC) and LUAD samples identified by scGPD (rows represent individual cells and columns represent genes, grouped by sample condition). **B**. UMAP projections of single-cell transcriptomes colored by gene expression and sample condition. **C**. GO enrichment analysis of scGPD identified genes.

The UMAP plot in the bottom right of Figure 5B shows the distribution of diseased conditions. The remaining UMAP projections in Figure 5B and Supplementary Figure S3 illustrate the spatial distribution of these genes across different cell clusters, demonstrating their selective enrichment in specific LUAD subpopulations. These complementary visualizations collectively demonstrate the ability of scGPD to robustly classify disease-associated genes.

Our results demonstrate that scGPD could successfully identify genes with significantly higher expression in LUAD-associated cell types compared to their healthy counterparts, as visualized in Figure 5B. These expression patterns confirm the genes’ specificity to the tumor microenvironment and suggest their potential roles in LUAD development and progression. To understand the biological significance of these identified genes, we performed GO enrichment analysis on the scGPD-selected gene panel. The analysis reveals that these genes are enriched in tumor-specific pathways and functionally associated with oncogenic signaling cascades, immune evasion mechanisms, and metastatic progression (Figure 5C). This functional enrichment further validates the biological relevance of our gene selection approach.

Beyond gene identification, scGPD offers a robust framework for biomarker discovery and precision medicine applications. The genes identified through scGPD provide valuable candidate targets for early diagnosis, patient stratification, and therapeutic intervention. Moreover, additional genes identified by scGPD, which are not shown here, can be found in Supplementary Figure S6 providing an expanded repository of potential LUAD biomarkers and therapeutic targets.

These findings emphasize the critical role of scGPD can play in discovering disease-associated genes and advancing our understanding of LUAD pathogenesis. By efficiently identifying key molecular drivers, scGPD enables deeper insights into the complex transcriptional landscape of cancer biology.

## Discussion

We have introduced scGPD, a deep learning framework for gene panel design in targeted spatial transcriptomics using single-cell RNA sequencing as a reference. It addresses two challenges: selecting an informative gene set within limited panel size to capture cellular and spatial diversity, and accounting for correlated gene expression that complicates gene prioritization. By incorporating a versatile learning framework and a correlation-aware gating mechanism, scGPD learns to identify compact, non-redundant gene sets that retain high predictive power for downstream biological tasks.

Through benchmarking on multiple single-cell datasets, scGPD shows strong performance across different panel sizes and task settings. Notably, genes selected from single-cell data transfer well to spatial contexts, capturing coherent spatial expression patterns. This shows that scGPD can extract signals that are biologically meaningful and spatially informative.

A key feature of scGPD is its explicit modeling of gene–gene correlations during the feature selection process. In contrast to many existing methods that treat genes as independent features, scGPD uses a gating mechanism to discourage the simultaneous selection of highly correlated genes. This reduces redundancy and ensures that selected genes span diverse biological programs, leading to more efficient and informative panels. Such consideration is especially important for technologies like FISH-based spatial methods, where the number of measurable targets is constrained and each gene must add unique value.

Beyond unsupervised applications, scGPD is also adaptable to task-specific goals through supervised training. By modifying the objective function, the model can prioritize genes predictive of particular phenotypes, such as disease states or functional cell properties. This makes scGPD well-suited for translational and clinical applications, such as identifying disease markers for spatial pathology or constructing custom panels.

While our results demonstrate the practical utility of scGPD, there are important directions for future work. Experimental validation of scGPD designed panels in spatial profiling platforms, particularly multiplexed FISH or hybridization-based methods, will be essential to confirm the effectiveness of selected panels in real tissues [43]. Incorporating spatial priors into the gene selection process may further enhance panel informativeness and generalizability, particularly for identifying complex tissue architectures and microenvironments [44] [45]. Furthermore, extending the framework to integrate multiomics data, such as CITE-seq or scATAC-seq, could provide a more comprehensive view of cellular states and improve the robustness of gene panels across diverse experimental settings [46]. Together, these directions hold promise for further enhancing the applicability and impact of scGPD in spatial and single-cell genomics.

In conclusion, scGPD provides an interpretable and flexible framework for gene panel design that bridges the gap between single-cell and spatial transcriptomics. By leveraging deep learning to model complex dependencies in gene expression and allowing for adaptable design criteria, scGPD offers a robust solution to a fundamental bottleneck in spatial transcriptomics workflows. We anticipate that scGPD will be a valuable tool for both basic and translational research, enabling efficient and targeted exploration of tissue architecture, cellular organization, and disease mechanisms.

## Methods

scGPD is a deep learning framework that uses reference single-cell RNA-seq data to build gene panels tailored for specific experimental goals. The method identifies a small set of genes that accurately capture transcriptome-wide expression patterns. This is achieved through a specialized architecture that incorporates differentiable feature selection layers, allowing joint optimization of gene selection and downstream tasks via stochastic gradient descent [47].

The framework operates through two consecutive stages. In the first stage, scGPD reconstructs transcriptome-wide expression profiles using a subset of candidate genes. Instead of examining genes individually, the model uses a correlation-aware binary gating mechanism that identifies interdependent gene groups through a Gaussian copula with Binary Concrete distribution to model gene relationships [17]. Through the incorporation of a regularization strategy, the model effectively enforces sparsity and prioritizes the selection of the most informative gene within each correlated group. To properly match the statistical properties of scRNA-seq data, scGPD uses Poisson loss as its objective function [19, 20, 21, 22], which naturally handles the count-based nature of single-cell expression data. In the second stage, scGPD performs final gene selection by choosing exactly *k* genes from the reduced candidate set (of size *d*) obtained from the first stage. This selection process again uses a binary gating mechanism, but now applies masks based on Concrete distributions [18] to enforce a strict limit of exactly *k* genes. To ensure the final gene panel works well for specific biological tasks, the model uses task-specific loss functions designed for target applications such as cell-type classification.

### Poisson loss function

Poisson loss is a suitable choice for modeling scRNA-seq data due to the distributional properties of gene expression counts [19, 20, 21, 22]. This characteristic makes Poisson loss particularly well suited for predicting transcriptome-wide expression from a selected subset of genes, as it captures the inherent statistical properties of scRNA-seq data while remaining computationally efficient.

Given observed gene expression counts *y*_*i*_ and predicted values *ŷ*_*i*_, the Poisson loss is defined as

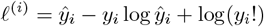

In scGPD, Poisson loss is used as the default objective in the first training stage, where the goal is to reconstruct transcriptome-wide expression and reduce the influence of uninformative genes. This loss function is well suited for modeling the count-based structure of scRNA-seq data, encouraging the model to prioritize genes that provide informative signals for transcriptome reconstruction. Poisson loss can incorporate cell-specific size factors, allowing the model to account for sequencing depth differences, thereby improving robustness to varying data coverage. While Poisson loss is the default, other loss functions such as mean squared error (MSE) can also be used, depending on the specific characteristics of the dataset or the downstream application. This flexibility allows scGPD to adapt to a variety of modeling needs while maintaining its focus on compact, informative gene panel design.

### Feature selection layers

Our methodology implements a two-stage feature selection framework. In the first stage, a correlated binary gating mechanism is used to filter out redundant and uninformative genetic markers. This design directs computational resources toward salient features by applying learned correlated binary gates (Supplementary Simulation Study, Figure S4). Rather than presuming selection independence, we characterize the gate vector distribution *B*_*i*_ using a correlated binary concrete formulation, expressed as *B*_*i*_ ∼ BinConcrete(*β*_*i*_, *τ*), where *β*_*i*_ denotes non-normalized logarithmic probabilities [18]. These probabilities are formulated via a Gaussian copula [17], introducing interdependencies between binary variables.

A Gaussian copula constitutes a multivariate cumulative distribution function for random variables *U*_1_, …, *U*_*p*_, defined across the unit hypercube [0, 1]^*p*^ with uniform marginal distributions where *U*_*k*_ ∼ Uniform(0, 1) for all *k* ∈ [*p*]. Given correlation matrix *R* ∈ [−1, 1]^*p*^ encoding the feature correlation structure of *X*, the Gaussian copula is formally defined as:

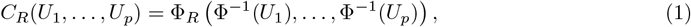

where Φ_*R*_ represents the joint cumulative distribution function of a multivariate Gaussian with zero mean and correlation matrix *R*, while Φ^−1^ denotes the inverse cumulative distribution function of a standard univariate Gaussian.

The logarithmic probabilities *β*_*i*_ are computed according to:

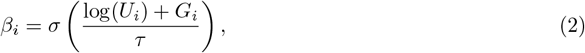

where *σ*(*x*) represents the sigmoid activation function, and *τ* constitutes the temperature hyperparameter. As *τ* approaches zero, the distribution converges to a Bernoulli distribution, whereas increasing *τ* toward infinity renders the distribution increasingly continuous.

The utilization of a Gaussian copula ensures that the gate vector distribution *B*_*i*_ accurately captures the correlation structure among input features, enabling the model to account for gate interdependencies during genetic marker selection.

The output from the correlated binary gates is determined by the Hadamard product **b** ⊙ **x**, where **x** represents the input vector and **b** = [*b*_1_, …, *b*_*p*_] ∈ ℝ^*p*^ comprises samples *b*_*i*_ drawn from the respective random variables *B*_*i*_ ∼ BinConcrete(*β*_*i*_, *τ*). This product subsequently traverses a neural architecture *f*_*θ*_ to predict gene expression counts **y**. Genetic marker elimination occurs when the model converges toward low values of *β*_*i*_, a process incentivized through the incorporation of a regularization term on BinConcrete samples within the objective function:

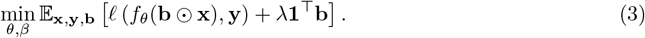

The regularization component *λ***1**^⊤^**b** penalizes excessive feature selection, with hyperparameter *λ >* 0 acts as a tuning parameter that balances predictive accuracy against feature sparsity, while *𝓁* denotes the Poisson loss function.

In the second stage, after constraining the candidate pool to size *d*, we again apply a binary mask to select exactly *k* genetic markers from the *d* candidates. The mask is generated via the element-wise maximum of *k* Concrete random variables, denoted as **A**_*i*_ ∼ Concrete(*α*_*i*_, *τ*) for *i* = 1, …, *k*. Each Concrete distribution is parameterized by non-normalized probabilities 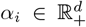 and temperature parameter *τ >* 0, with each distribution asymptotically approaching a multinomial distribution with probabilities defined by 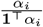 as *τ* approaches zero [18].

The binary mask layer processes normalized gene expression profiles **x** ∈ ℝ^*d*^, yielding the Hadamard product **a**⊙**x**, where **a** = max_*i*_ **a**_*i*_ ∈ ℝ^*d*^ constitutes the element-wise maximum of samples **a**_*i*_ drawn from each concrete distribution random variable. This product subsequently traverses a neural architecture *f*_*γ*_, which predicts the target variable **y** based on **a** ⊙ **x**. The model parameters are optimized according to:

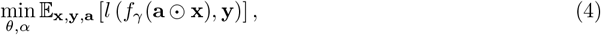

where *l* represents an application-specific loss function.

## Supporting information

Supplementary information for scGPD

## Acknowledgments

This work was supported in part by NIH grants U01 HG013840 and U24 HG012108.

## Data availability

We provide a summary of the sources and statistics for all datasets used in Supplementary Table S5. All datasets are accessible through the links included in this file.

## Code availability

The codes of scGPD are available at https://github.com/TinaGuo/scGPD. We follow the MIT license for usage.

## References

[1] Dragomirka Jovic, Xue Liang, Hua Zeng, et al. Single-cell RNA sequencing technologies and applications: A brief overview. Clinical and translational medicine 12.3 (2022), e694.

[2] Evan Z Macosko, Anindita Basu, Rahul Satija, et al. Highly parallel genome-wide expression profiling of individual cells using nanoliter droplets. Cell 161.5 (2015), pp. 1202–1214.

[3] Fuchou Tang, Catalin Barbacioru, Yangzhou Wang, et al. mRNA-Seq whole-transcriptome analysis of a single cell. Nature methods 6.5 (2009), pp. 377–382.

[4] Grace XY Zheng, Jessica M Terry, Phillip Belgrader, et al. Massively parallel digital transcriptional profiling of single cells. Nature communications 8.1 (2017), p. 14049.

[5] Jun Du, Yu-Chen Yang, Zhi-Jie An, et al. Advances in spatial transcriptomics and related data analysis strategies. Journal of translational medicine 21.1 (2023), p. 330.

[6] Guohao Liang, Hong Yin, and Fangyuan Ding. Technical advances and applications of spatial transcriptomics. GEN biotechnology 2.5 (2023), pp. 384–398.

[7] Andrea M Femino, Fredric S Fay, Kevin Fogarty, et al. Visualization of single RNA transcripts in situ. Science 280.5363 (1998), pp. 585–590.

[8] Simone Codeluppi, Lars E Borm, Amit Zeisel, et al. Spatial organization of the somatosensory cortex revealed by osmFISH. Nature methods 15.11 (2018), pp. 932–935.

[9] Kok Hao Chen, Alistair N Boettiger, Jeffrey R Moffitt, et al. Spatially resolved, highly multiplexed RNA profiling in single cells. Science 348.6233 (2015), aaa6090.

[10] 10x Genomics. Visium Spatial Gene Expression. Available at: https://www.10xgenomics.com/spatial-transcriptomics. 2020.

[11] Sanja Vickovic, Gökcen Eraslan, Fredrik Salmén, et al. High-definition spatial transcriptomics for in situ tissue profiling. Nature methods 16.10 (2019), pp. 987–990.

[12] Chee-Huat Linus Eng, Michael Lawson, Qian Zhu, et al. Transcriptome-scale super-resolved imaging in tissues by RNA seqFISH+. Nature 568.7751 (2019), pp. 235–239.

[13] Il-Gyo Chong and Chi-Hyuck Jun. Performance of some variable selection methods when multicollinearity is present. Chemometrics and intelligent laboratory systems 78.1-2 (2005), pp. 103–112.

[14] Alexandr Katrutsa and Vadim Strijov. Comprehensive study of feature selection methods to solve multicollinearity problem according to evaluation criteria. Expert Systems with Applications 76 (2017), pp. 1–11.

[15] David A Belsley, Edwin Kuh, and Roy E Welsch. Regression diagnostics: Identifying influential data and sources of collinearity. John Wiley & Sons, 2005.

[16] Changhee Lee, Fergus Imrie, and Mihaela van der Schaar. Self-supervision enhanced feature selection with correlated gates. International conference on learning representations. 2022.

[17] Roger B Nelsen. An introduction to copulas. Springer, 2006.

[18] Chris J Maddison, Andriy Mnih, and Yee Whye Teh. The concrete distribution: A continuous relaxation of discrete random variables. arXiv preprint arXiv:1611.00712 (2016).

[19] Shiquan Sun, Jiaqiang Zhu, and Xiang Zhou. Statistical analysis of spatial expression patterns for spatially resolved transcriptomic studies. Nature methods 17.2 (2020), pp. 193–200.

[20] Dylan M Cable, Evan Murray, Luli S Zou, et al. Robust decomposition of cell type mixtures in spatial transcriptomics. Nature biotechnology 40.4 (2022), pp. 517–526.

[21] Dylan M Cable, Evan Murray, Vignesh Shanmugam, et al. Cell type-specific inference of differential expression in spatial transcriptomics. Nature methods 19.9 (2022), pp. 1076–1087.

[22] Gefei Wang, Jia Zhao, Yan Yan, et al. Construction of a 3D whole organism spatial atlas by joint modelling of multiple slices with deep neural networks. Nature Machine Intelligence 5.11 (2023), pp. 1200–1213.

[23] Maayan Baron, Adrian Veres, Samuel L Wolock, et al. A single-cell transcriptomic map of the human and mouse pancreas reveals inter-and intra-cell population structure. Cell systems 3.4 (2016), pp. 346–360.

[24] Monika Litviňuková, Carlos Talavera-López, Henrike Maatz, et al. Cells of the adult human heart. Nature 588.7838 (2020), pp. 466–472.

[25] Chang Su, Zichun Xu, Xinning Shan, et al. Cell-type-specific co-expression inference from single cell RNA-sequencing data. Nature Communications 14.1 (2023), p. 4846.

[26] Ian Covert, Rohan Gala, Tim Wang, et al. Predictive and robust gene selection for spatial transcriptomics. Nature Communications 14.1 (2023), p. 2091.

[27] Mashrur Ahmed Yafi, Md Hasibul Husain Hisham, Francisco Grisanti, et al. scGIST: gene panel design for spatial transcriptomics with prioritized gene sets. Genome Biology 25.1 (2024), p. 57.

[28] Alsu Missarova, Jaison Jain, Andrew Butler, et al. geneBasis: an iterative approach for unsupervised selection of targeted gene panels from scRNA-seq. Genome biology 22 (2021), pp. 1–22.

[29] Philip Sedgwick. Pearson’s correlation coefficient. Bmj 345 (2012).

[30] David H Brann, Tatsuya Tsukahara, Caleb Weinreb, et al. Non-neuronal expression of SARS-CoV-2 entry genes in the olfactory system suggests mechanisms underlying COVID-19-associated anosmia. Science advances 6.31 (2020), eabc5801.

[31] F Alexander Wolf, Fiona K Hamey, Mireya Plass, et al. PAGA: graph abstraction reconciles clustering with trajectory inference through a topology preserving map of single cells. Genome biology 20 (2019), pp. 1–9.

[32] Sunny Z Wu, Ghamdan Al-Eryani, Daniel Lee Roden, et al. A single-cell and spatially resolved atlas of human breast cancers. Nature genetics 53.9 (2021), pp. 1334–1347.

[33] Bianca Pellegrino, Keren David, Stav Rabani, et al. CD74 promotes the formation of an immunosuppressive tumor microenvironment in triple-negative breast cancer in mice by inducing the expansion of tolerogenic dendritic cells and regulatory B cells. PLoS Biology 22.11 (2024), e3002905.

[34] Xiang Li, Zeng Li, Shanzhi Gu, et al. A pan-cancer analysis of collagen VI family on prognosis, tumor microenvironment, and its potential therapeutic effect. BMC bioinformatics 23.1 (2022), p. 390.

[35] Ping Yin, Yu Bai, Zhuo Wang, et al. Non-canonical Fzd7 signaling contributes to breast cancer mesenchymal-like stemness involving Col6a1. Cell Communication and Signaling 18 (2020), pp. 1–13.

[36] Nayoung Kim, Hong Kwan Kim, Kyungjong Lee, et al. Single-cell RNA sequencing demonstrates the molecular and cellular reprogramming of metastatic lung adenocarcinoma. Nature communications 11.1 (2020), p. 2285.

[37] David J Myers and Jason M Wallen. Lung adenocarcinoma. StatPearls [Internet]. StatPearls Publishing, 2023.

[38] Huaqiang Zhou, Yaxiong Zhang, Jiaqing Liu, et al. Education and lung cancer: a Mendelian randomization study. International journal of epidemiology 48.3 (2019), pp. 743–750.

[39] Xiwei Sun, Jiani Yi, Juze Yang, et al. An integrated epigenomic-transcriptomic landscape of lung cancer reveals novel methylation driver genes of diagnostic and therapeutic relevance. Theranostics 11.11 (2021), p. 5346.

[40] Hongxia Tian, Kezhen Li, Jing Chen, et al. Impaired natural killer cell maturation in lung adenocarcinoma driven by FABP4 and SPON2 downregulation through disrupted lipid metabolism. Translational Lung Cancer Research 14.5 (2025), p. 1660.

[41] MJ Duffy and K McCarthy. Matrix metalloproteinases in cancer: prognostic markers and targets for therapy. International journal of oncology 12.6 (1998), pp. 1343–1351.

[42] Srinivas Mamidi, Simon Höne, and Michael Kirschfink. The complement system in cancer: Ambivalence between tumour destruction and promotion. Immunobiology 222.1 (2017), pp. 45–54.

[43] Christopher R Merritt, Giang T Ong, Sarah E Church, et al. Multiplex digital spatial profiling of proteins and RNA in fixed tissue. Nature biotechnology 38.5 (2020), pp. 586–599.

[44] Rohit Arora, Christian Cao, Mehul Kumar, et al. Spatial transcriptomics reveals distinct and conserved tumor core and edge architectures that predict survival and targeted therapy response. Nature Communications 14.1 (2023), p. 5029.

[45] Vitalii Kleshchevnikov, Artem Shmatko, Emma Dann, et al. Cell2location maps fine-grained cell types in spatial transcriptomics. Nature biotechnology 40.5 (2022), pp. 661–671.

[46] Eleni P Mimitou, Caleb A Lareau, Kelvin Y Chen, et al. Scalable, multimodal profiling of chromatin accessibility, gene expression and protein levels in single cells. Nature biotechnology 39.10 (2021), pp. 1246–1258.

[47] Shun-ichi Amari. Backpropagation and stochastic gradient descent method. Neurocomputing 5.4-5 (1993), pp. 185–196.

