## Supplementary information for scGPD for "scGPD: single-cell informed gene panel design for targeted spatial transcriptomics"

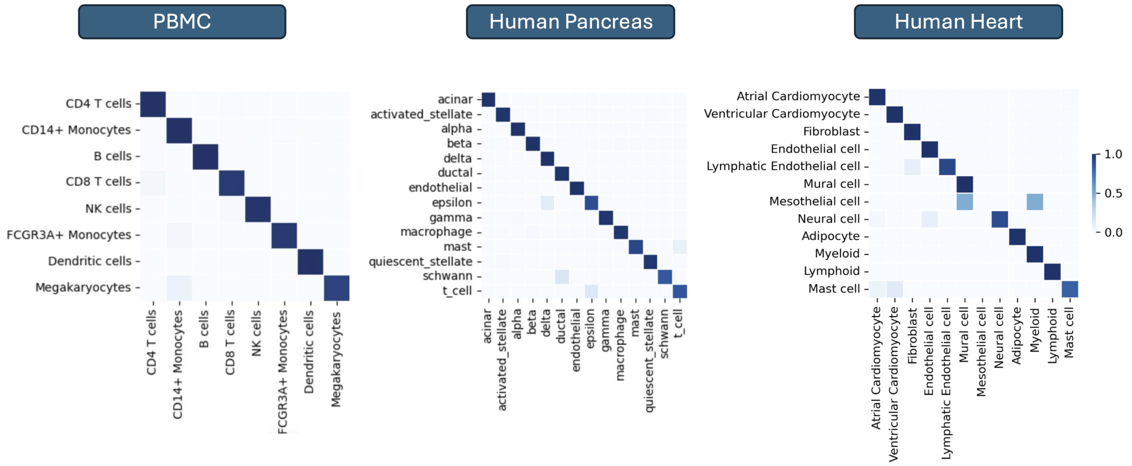


Figure S1: Confusion matrix of dataset PBMC, Human pancreas and Human heart for gene size of 256.

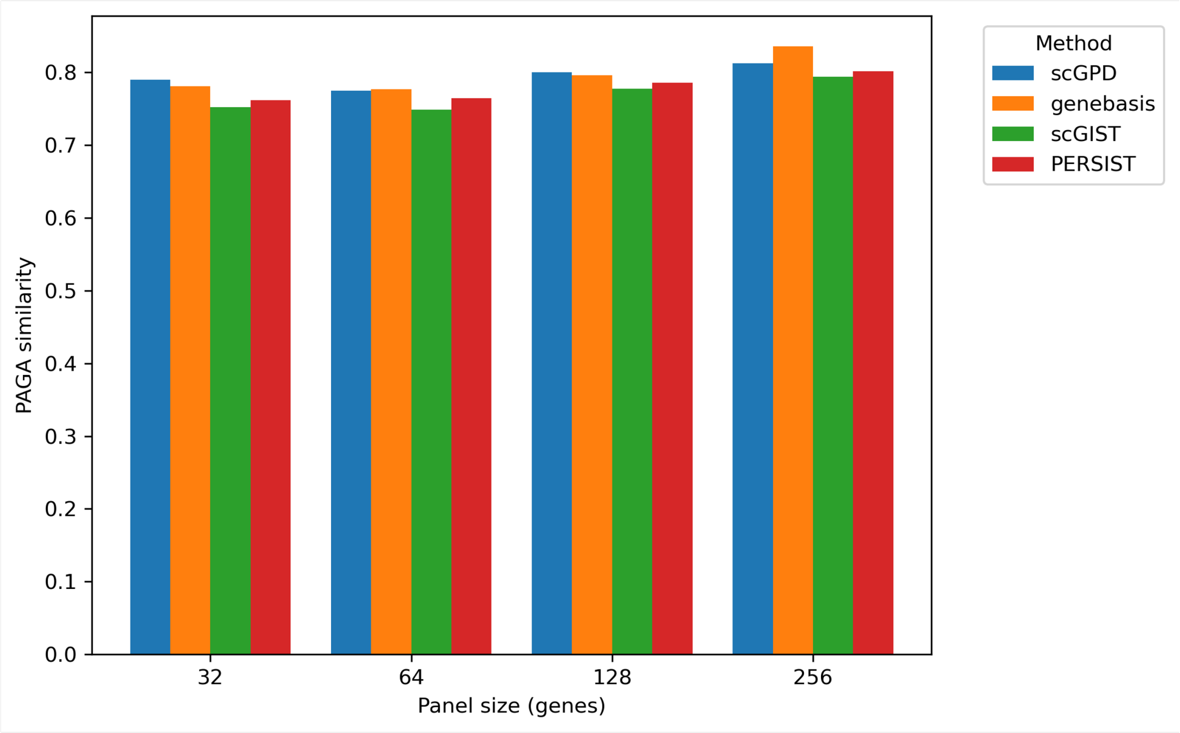


Figure S2: PAGA similarity scores across different panel sizes for different methods, reflecting the ability of each method to preserve the underlying cellular topology of the seqFISH+ dataset.

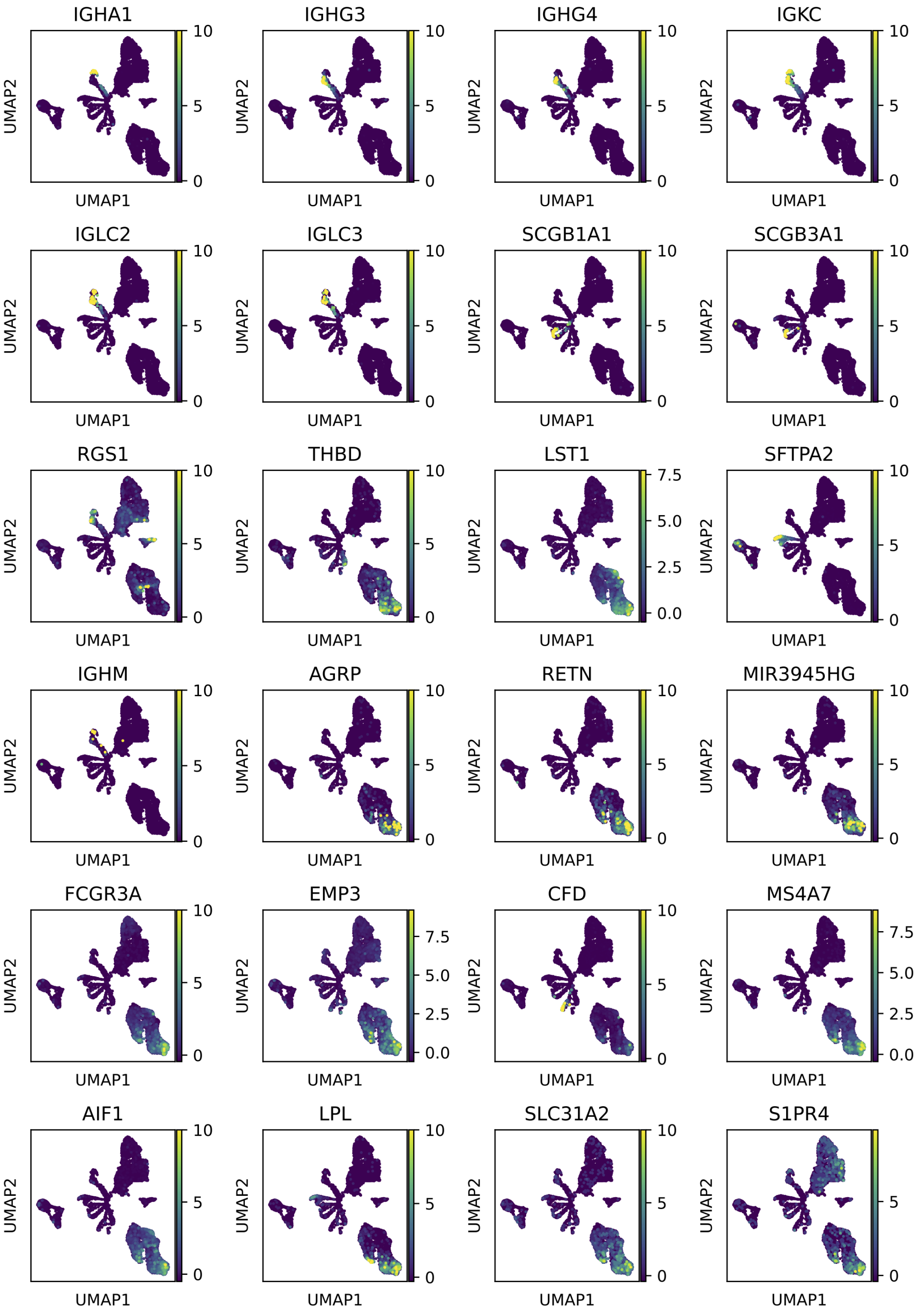
IGHA1
IGHG3
IGHG4
IGKC
IGLC2
IGLC3
SCGB1A1
SCGB3A1
RGS1
THBD
LST1
SFTPA2
IGHM
AGRP
RETN
MIR3945HG
FCGR3A
EMP3
CFD
MS4A7
AIF1
LPL
SLC31A2
S1PR4
UMAP1
UMAP2

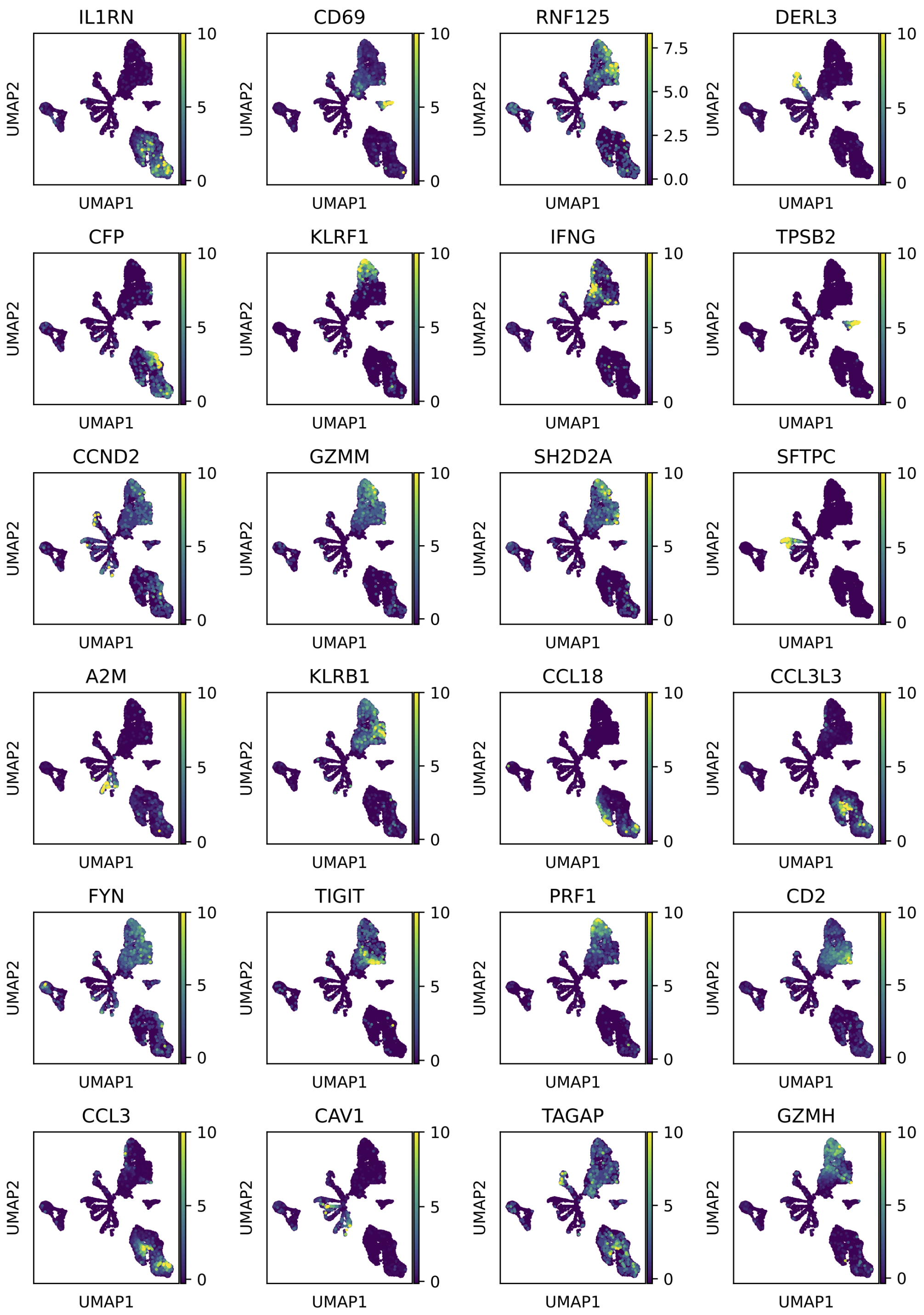


Figure S3: UMAP projections of the LUAD-related genes and their expression dynamics across cell types.

| Table S1. p-values of 32 identified genes from human brest cancer dataset | | |
|---|---|---|
| Gene | combined p-value | adjusted p-value |
| CXCL14 | 3.54E-259 | 4.59E-257 |
| KRT19 | 1.06E-194 | 6.87E-193 |
| CEACAM6 | 2.29E-178 | 9.90E-177 |
| TIMP2 | 2.79E-166 | 9.07E-165 |
| SPARC | 2.74E-158 | 7.10E-157 |
| CD24 | 2.24E-154 | 4.86E-153 |
| KRT7 | 9.37E-124 | 1.74E-122 |
| COL1A1 | 7.45E-114 | 1.21E-112 |
| BGN | 1.93E-90 | 2.78E-89 |
| SCD | 2.22E-74 | 2.88E-73 |
| HLA-DRA | 1.07E-72 | 1.27E-71 |
| SERPING1 | 8.19E-72 | 8.87E-71 |
| SERPINF1 | 6.17E-63 | 6.17E-62 |
| LRP1 | 1.94E-60 | 1.80E-59 |
| RARRES2 | 2.17E-55 | 1.86E-54 |
| PRLR | 2.30E-55 | 1.86E-54 |
| VCAN | 5.53E-51 | 4.23E-50 |
| GPR160 | 3.56E-47 | 2.57E-46 |
| TTC39A | 8.50E-39 | 5.81E-38 |
| MGST1 | 1.05E-31 | 6.81E-31 |
| NUPR2 | 1.12E-26 | 6.94E-26 |
| ANGPTL2 | 4.06E-25 | 2.40E-24 |
| ELN | 3.67E-23 | 2.07E-22 |
| CXCL12 | 8.59E-21 | 4.65E-20 |
| CD79A | 9.22E-20 | 4.79E-19 |
| CYP1B1 | 1.34E-18 | 6.68E-18 |
| PSCA | 6.80E-17 | 3.27E-16 |
| CRISPLD2 | 2.35E-16 | 1.09E-15 |
| IGF1 | 2.54E-09 | 1.14E-08 |
| FCRL5 | 1.16E-07 | 5.03E-07 |
| GAPT | 1.09E-04 | 4.55E-04 |
| ARHGAP24 | 2.19E-03 | 8.87E-03 |

Table S2. p-values of 64 identified genes from human brest cancer dataset

| Gene | combined p-value | adjusted p-value |
|---|---|---|
| ELF3 | 5.76E-270 | 1.68E-267 |
| KRT19 | 1.06E-194 | 1.55E-192 |
| FN1 | 6.92E-179 | 6.74E-177 |
| TIMP2 | 2.79E-166 | 2.04E-164 |
| SPARC | 2.74E-158 | 1.60E-156 |
| PITX1 | 9.88E-149 | 4.81E-147 |
| CD74 | 1.10E-139 | 4.59E-138 |
| KRT7 | 9.37E-124 | 3.42E-122 |
| AZGP1 | 1.29E-117 | 4.19E-116 |
| TNNT1 | 3.26E-116 | 9.53E-115 |
| COLEC12 | 1.36E-114 | 3.62E-113 |
| COL6A1 | 5.33E-111 | 1.30E-109 |
| IRX3 | 7.29E-103 | 1.64E-101 |
| LUM | 2.22E-101 | 4.64E-100 |
| RAC3 | 4.84E-96 | 9.42E-95 |
| CTHRC1 | 3.86E-95 | 7.05E-94 |
| ISLR | 6.35E-93 | 1.09E-91 |
| ST8SIA6.AS1 | 6.84E-87 | 1.11E-85 |
| MFAP2 | 2.37E-85 | 3.65E-84 |
| MAL2 | 3.90E-84 | 5.70E-83 |
| LMTK3 | 4.00E-77 | 5.57E-76 |
| SERPING1 | 8.19E-72 | 1.09E-70 |
| EPCAM | 4.91E-65 | 6.23E-64 |
| SERPINF1 | 6.17E-63 | 7.51E-62 |
| LRP1 | 1.94E-60 | 2.27E-59 |
| ITGA11 | 2.48E-59 | 2.79E-58 |
| IGFBP4 | 5.01E-58 | 5.42E-57 |
| IGFBP7 | 1.86E-57 | 1.94E-56 |
| RARRES2 | 2.17E-55 | 2.19E-54 |
| PCOLCE | 9.15E-55 | 8.91E-54 |
| S100A1 | 4.36E-54 | 4.11E-53 |
| INHBA | 1.60E-51 | 1.46E-50 |
| CALD1 | 5.81E-51 | 5.14E-50 |
| CTGF | 1.36E-47 | 1.17E-46 |
| C3 | 2.10E-46 | 1.75E-45 |
| CA12 | 1.30E-42 | 1.06E-41 |
| DNAJC22 | 2.38E-38 | 1.88E-37 |
| SRPX2 | 1.98E-37 | 1.53E-36 |
| TNFAIP6 | 5.03E-35 | 3.77E-34 |
| RCN3 | 1.47E-33 | 1.08E-32 |
| HOXC10 | 6.30E-32 | 4.49E-31 |
| MGST1 | 1.05E-31 | 7.29E-31 |
| OVOL1 | 5.70E-21 | 3.88E-20 |
| TIMP1 | 6.96E-20 | 4.62E-19 |

| | | |
|---|---|---|
| CD79A | 9.22E-20 | 5.99E-19 |
| MS4A1 | 9.98E-17 | 6.34E-16 |
| LAMP5 | 4.11E-15 | 2.55E-14 |
| S100A7 | 7.66E-11 | 4.66E-10 |
| TMSB15A | 6.89E-10 | 4.11E-09 |
| IGF1 | 2.54E-09 | 1.48E-08 |
| ITGBL1 | 3.19E-09 | 1.83E-08 |
| IGHM | 5.76E-09 | 3.24E-08 |
| VPREB3 | 7.76E-08 | 4.28E-07 |
| FCRL5 | 1.16E-07 | 6.29E-07 |
| ADAM28 | 2.20E-06 | 1.17E-05 |
| PNOC | 1.02E-04 | 5.33E-04 |
| SVEP1 | 1.64E-04 | 8.39E-04 |
| CCDC8 | 2.41E-04 | 1.21E-03 |
| RUNX1T1 | 2.54E-04 | 1.26E-03 |
| RGMA | 3.02E-02 | 1.47E-01 |
| TCEAL5 | 6.83E-01 | 1.00E+00 |
| TFAP2B | 3.76E-01 | 1.00E+00 |

| Table S3. p-values of 128 identified genes from human brest cancer dataset | | |
|---|---|---|
| Gene | combined p-value | adjusted p-value |
| KRT8 | 0.00E+00 | 0.00E+00 |
| FXYD3 | 0.00E+00 | 0.00E+00 |
| DEGS2 | 0.00E+00 | 0.00E+00 |
| SMIM22 | 4.23E-295 | 7.28E-293 |
| CALML5 | 1.57E-276 | 2.16E-274 |
| CXCL14 | 3.54E-259 | 4.06E-257 |
| AGR2 | 6.26E-255 | 6.16E-253 |
| CLDN4 | 1.45E-204 | 1.25E-202 |
| FN1 | 6.92E-179 | 5.30E-177 |
| CEACAM6 | 2.29E-178 | 1.58E-176 |
| TIMP2 | 2.79E-166 | 1.75E-164 |
| S100P | 7.92E-162 | 4.55E-160 |
| DHCR24 | 2.44E-158 | 1.29E-156 |
| SPINT2 | 3.36E-158 | 1.65E-156 |
| POSTN | 2.34E-150 | 1.07E-148 |
| FASN | 4.20E-150 | 1.81E-148 |
| CD74 | 1.10E-139 | 4.46E-138 |
| AR | 3.50E-139 | 1.34E-137 |
| AZGP1 | 1.29E-117 | 4.68E-116 |
| S100A14 | 1.09E-116 | 3.74E-115 |
| TNNT1 | 3.26E-116 | 1.07E-114 |
| COLEC12 | 1.36E-114 | 4.26E-113 |
| RAB25 | 1.75E-112 | 5.23E-111 |
| CLDN7 | 1.43E-110 | 4.11E-109 |
| ABCC11 | 1.11E-97 | 3.05E-96 |
| RAC3 | 4.84E-96 | 1.28E-94 |
| ANTXR1 | 8.02E-96 | 2.05E-94 |
| CTHRC1 | 3.86E-95 | 9.50E-94 |
| BGN | 1.93E-90 | 4.58E-89 |
| DSCAM.AS1 | 6.68E-89 | 1.53E-87 |
| ST8SIA6.AS1 | 6.84E-87 | 1.52E-85 |
| MFAP2 | 2.37E-85 | 5.11E-84 |
| CDH11 | 6.51E-74 | 1.36E-72 |
| HLA.DRA | 1.07E-72 | 2.18E-71 |
| SERPING1 | 8.19E-72 | 1.61E-70 |
| F12 | 1.15E-69 | 2.20E-68 |
| CST1 | 9.31E-68 | 1.73E-66 |
| SERPINF1 | 6.17E-63 | 1.12E-61 |
| MESP1 | 2.94E-62 | 5.19E-61 |
| LRP1 | 1.94E-60 | 3.34E-59 |
| SFRP2 | 8.17E-59 | 1.37E-57 |
| IGFBP4 | 5.01E-58 | 8.21E-57 |
| IGFBP7 | 1.86E-57 | 2.99E-56 |
| HOXC11 | 5.08E-57 | 7.96E-56 |

| | | |
|---|---|---|
| TMC5 | 5.09E-55 | 7.79E-54 |
| PCOLCE | 9.15E-55 | 1.37E-53 |
| S100A1 | 4.36E-54 | 6.40E-53 |
| VCAN | 5.53E-51 | 7.94E-50 |
| CALD1 | 5.81E-51 | 8.17E-50 |
| AARD | 7.85E-51 | 1.08E-49 |
| CTGF | 1.36E-47 | 1.84E-46 |
| GPR160 | 3.56E-47 | 4.72E-46 |
| ALDH3B2 | 3.94E-46 | 5.12E-45 |
| TOM1L1 | 1.11E-43 | 1.41E-42 |
| DPYSL3 | 1.17E-43 | 1.47E-42 |
| SPON2 | 3.16E-43 | 3.88E-42 |
| MXRA8 | 2.15E-42 | 2.60E-41 |
| CLEC11A | 1.04E-39 | 1.24E-38 |
| EFNA3 | 3.27E-39 | 3.82E-38 |
| TTC39A | 8.50E-39 | 9.76E-38 |
| FSTL1 | 1.08E-38 | 1.21E-37 |
| NID1 | 2.99E-38 | 3.32E-37 |
| SRPX2 | 1.98E-37 | 2.17E-36 |
| TNFAIP6 | 5.03E-35 | 5.41E-34 |
| MGST1 | 1.05E-31 | 1.11E-30 |
| SCGB2A2 | 2.91E-30 | 3.04E-29 |
| MFAP5 | 1.74E-29 | 1.79E-28 |
| C1S | 2.28E-29 | 2.31E-28 |
| CCDC80 | 1.25E-28 | 1.25E-27 |
| NUPR2 | 1.12E-26 | 1.11E-25 |
| ANGPTL2 | 4.06E-25 | 3.94E-24 |
| FBLN1 | 4.09E-24 | 3.92E-23 |
| ELN | 3.67E-23 | 3.46E-22 |
| FBLN2 | 4.28E-22 | 3.98E-21 |
| NID2 | 4.84E-22 | 4.45E-21 |
| FSIP1 | 1.26E-21 | 1.14E-20 |
| CXCL12 | 8.59E-21 | 7.69E-20 |
| CD79A | 9.22E-20 | 8.14E-19 |
| FIBIN | 2.04E-19 | 1.78E-18 |
| SSC5D | 2.75E-19 | 2.37E-18 |
| EFEMP1 | 4.39E-19 | 3.74E-18 |
| PDGFRL | 2.50E-18 | 2.10E-17 |
| GRP | 7.05E-18 | 5.85E-17 |
| MS4A1 | 9.98E-17 | 8.18E-16 |
| CD79B | 1.35E-14 | 1.10E-13 |
| ATP2C2 | 2.75E-13 | 2.21E-12 |
| TNFRSF13C | 2.84E-13 | 2.25E-12 |
| MSX2 | 2.79E-12 | 2.18E-11 |
| SPIB | 3.32E-12 | 2.57E-11 |
| GOLT1A | 5.64E-11 | 4.32E-10 |
| S100A7 | 7.66E-11 | 5.80E-10 |
| SOX11 | 1.39E-10 | 1.04E-09 |

| | | |
|---|---|---|
| CD19 | 3.19E-10 | 2.36E-09 |
| TMSB15A | 6.89E-10 | 5.05E-09 |
| FBLN5 | 1.25E-08 | 9.06E-08 |
| CLMP | 2.14E-08 | 1.54E-07 |
| HOXB13 | 3.14E-08 | 2.23E-07 |
| VPREB3 | 7.76E-08 | 5.46E-07 |
| FCRL5 | 1.16E-07 | 8.09E-07 |
| SDR16C5 | 1.01E-06 | 6.98E-06 |
| PNOC | 1.02E-04 | 6.96E-04 |
| GAPT | 1.09E-04 | 7.34E-04 |
| CCDC8 | 2.41E-04 | 1.61E-03 |
| BOC | 2.53E-04 | 1.68E-03 |
| TMEM40 | 5.62E-04 | 3.69E-03 |
| DCLK1 | 9.18E-04 | 5.97E-03 |
| ABCA8 | 1.34E-03 | 8.62E-03 |
| COL6A6 | 2.37E-03 | 1.51E-02 |
| IGHD | 3.30E-03 | 2.08E-02 |
| MEDAG | 3.45E-03 | 2.16E-02 |
| SLIT2 | 5.27E-03 | 3.27E-02 |
| ABCA9 | 6.00E-03 | 3.69E-02 |
| BMPR1B | 7.16E-03 | 4.36E-02 |
| PLA2G10 | 1.09E-02 | 6.59E-02 |
| PAPLN | 1.52E-02 | 9.13E-02 |
| LSAMP | 1.85E-02 | 1.10E-01 |
| LINC01140 | 8.83E-02 | 5.20E-01 |
| DKK1 | 1.54E-01 | 8.96E-01 |
| MEIS2 | 4.04E-01 | 1.00E+00 |
| MMP16 | 2.80E-01 | 1.00E+00 |
| HAND2.AS1 | 9.16E-01 | 1.00E+00 |
| RSPO1 | 6.56E-01 | 1.00E+00 |
| CACNA1G | 3.20E-01 | 1.00E+00 |
| TCEAL5 | 6.83E-01 | 1.00E+00 |
| LINC01133 | 7.00E-01 | 1.00E+00 |
| LINC01285 | 5.29E-01 | 1.00E+00 |
| MSLN | 3.85E-01 | 1.00E+00 |

| Table S4. p-values of 256 identified genes from human brest cancer dataset | | |
|---|---|---|
| Gene | combined p-value | adjusted p-value |
| KRT18 | 0.00E+00 | 0.00E+00 |
| FXYD3 | 0.00E+00 | 0.00E+00 |
| MUC1 | 0.00E+00 | 0.00E+00 |
| CALML5 | 1.57E-276 | 6.02E-274 |
| CLDN3 | 1.52E-272 | 4.69E-270 |
| CXCL14 | 3.54E-259 | 9.08E-257 |
| SERINC2 | 4.01E-247 | 8.81E-245 |
| AGR3 | 5.07E-237 | 9.75E-235 |
| C19orf33 | 5.17E-221 | 8.84E-219 |
| CLDN4 | 1.45E-204 | 2.23E-202 |
| KRT19 | 1.06E-194 | 1.48E-192 |
| LRRC26 | 7.68E-193 | 9.85E-191 |
| MISP | 4.84E-188 | 5.73E-186 |
| FN1 | 6.92E-179 | 7.61E-177 |
| CEACAM6 | 2.29E-178 | 2.35E-176 |
| SPDEF | 2.54E-173 | 2.45E-171 |
| AEBP1 | 8.82E-169 | 7.98E-167 |
| TIMP2 | 2.79E-166 | 2.39E-164 |
| PRSS8 | 1.98E-163 | 1.61E-161 |
| S100P | 7.92E-162 | 6.09E-160 |
| DHCR24 | 2.44E-158 | 1.79E-156 |
| SPARC | 2.74E-158 | 1.91E-156 |
| SPINT2 | 3.36E-158 | 2.25E-156 |
| CD24 | 2.24E-154 | 1.44E-152 |
| POSTN | 2.34E-150 | 1.44E-148 |
| PITX1 | 9.88E-149 | 5.85E-147 |
| CD74 | 1.10E-139 | 6.27E-138 |
| COL12A1 | 2.15E-130 | 1.18E-128 |
| SULT2B1 | 2.44E-130 | 1.30E-128 |
| CYP4B1 | 2.48E-129 | 1.27E-127 |
| SLC44A4 | 1.07E-127 | 5.31E-126 |
| IRX5 | 2.76E-124 | 1.33E-122 |
| KRT7 | 9.37E-124 | 4.37E-122 |
| COMP | 6.66E-118 | 3.01E-116 |
| HTRA1 | 1.86E-116 | 8.20E-115 |
| TNNT1 | 3.26E-116 | 1.40E-114 |
| COLEC12 | 1.36E-114 | 5.66E-113 |
| C2orf54 | 2.02E-114 | 8.19E-113 |
| COL1A1 | 7.45E-114 | 2.94E-112 |
| RAB25 | 1.75E-112 | 6.72E-111 |
| COL6A1 | 5.33E-111 | 2.00E-109 |
| CLDN7 | 1.43E-110 | 5.24E-109 |
| COL5A1 | 8.84E-107 | 3.16E-105 |
| CRB3 | 1.84E-106 | 6.43E-105 |

| | | |
|---|---|---|
| KIF26B | 1.99E-105 | 6.82E-104 |
| LUM | 2.22E-101 | 7.44E-100 |
| THBS2 | 5.28E-101 | 1.73E-99 |
| ABCC11 | 1.11E-97 | 3.55E-96 |
| CNTD2 | 3.62E-95 | 1.14E-93 |
| ISLR | 6.35E-93 | 1.96E-91 |
| BGN | 1.93E-90 | 5.82E-89 |
| MMP2 | 2.00E-90 | 5.94E-89 |
| KLHDC9 | 2.98E-90 | 8.66E-89 |
| AP1M2 | 1.27E-87 | 3.61E-86 |
| ST8SIA6.AS1 | 6.84E-87 | 1.91E-85 |
| MFAP2 | 2.37E-85 | 6.52E-84 |
| COL6A3 | 1.07E-83 | 2.90E-82 |
| MRC2 | 9.18E-78 | 2.44E-76 |
| MLPH | 1.06E-77 | 2.78E-76 |
| TFF3 | 1.78E-76 | 4.57E-75 |
| FBN1 | 1.26E-74 | 3.19E-73 |
| SCD | 2.22E-74 | 5.50E-73 |
| CDH11 | 6.51E-74 | 1.59E-72 |
| HLA.DRA | 1.07E-72 | 2.58E-71 |
| SERPING1 | 8.19E-72 | 1.94E-70 |
| F12 | 1.15E-69 | 2.68E-68 |
| EVPL | 1.20E-67 | 2.76E-66 |
| C1orf116 | 1.48E-66 | 3.35E-65 |
| HTRA3 | 5.04E-66 | 1.12E-64 |
| EPCAM | 4.91E-65 | 1.08E-63 |
| PRR15 | 1.77E-64 | 3.83E-63 |
| DIO1 | 2.14E-63 | 4.58E-62 |
| CTXN1 | 1.49E-62 | 3.14E-61 |
| LRP1 | 1.94E-60 | 4.03E-59 |
| SFRP2 | 8.17E-59 | 1.68E-57 |
| P3H3 | 2.41E-58 | 4.88E-57 |
| IGFBP4 | 5.01E-58 | 1.00E-56 |
| IGFBP7 | 1.86E-57 | 3.68E-56 |
| HOXC11 | 5.08E-57 | 9.90E-56 |
| TMC5 | 5.09E-55 | 9.79E-54 |
| RAB17 | 1.40E-54 | 2.65E-53 |
| TFAP2A | 4.40E-54 | 8.17E-53 |
| S100A1 | 4.36E-54 | 8.17E-53 |
| TOX3 | 6.47E-53 | 1.19E-51 |
| LOXL1 | 9.57E-52 | 1.73E-50 |
| COL8A1 | 3.16E-51 | 5.65E-50 |
| COL8A2 | 4.45E-51 | 7.88E-50 |
| VCAN | 5.53E-51 | 9.68E-50 |
| CALD1 | 5.81E-51 | 1.00E-49 |
| TFF1 | 5.28E-50 | 9.02E-49 |
| SCGB1D2 | 8.97E-49 | 1.52E-47 |
| CTGF | 1.36E-47 | 2.28E-46 |

| | | |
|---|---|---|
| GPR160 | 3.56E-47 | 5.90E-46 |
| C3 | 2.10E-46 | 3.44E-45 |
| TOM1L1 | 1.11E-43 | 1.80E-42 |
| DPYSL3 | 1.17E-43 | 1.88E-42 |
| SPON2 | 3.16E-43 | 5.01E-42 |
| ADAMTS2 | 7.25E-43 | 1.14E-41 |
| CA12 | 1.30E-42 | 2.03E-41 |
| TMEM125 | 3.41E-42 | 5.25E-41 |
| MUC5B | 2.96E-41 | 4.52E-40 |
| AKR7A3 | 3.12E-40 | 4.71E-39 |
| EFEMP2 | 3.97E-40 | 5.93E-39 |
| CLEC11A | 1.04E-39 | 1.54E-38 |
| ASPN | 8.13E-39 | 1.19E-37 |
| FSTL1 | 1.08E-38 | 1.56E-37 |
| DNAJC22 | 2.38E-38 | 3.43E-37 |
| NID1 | 2.99E-38 | 4.25E-37 |
| SRPX2 | 1.98E-37 | 2.80E-36 |
| C1QTNF3 | 5.41E-37 | 7.56E-36 |
| COL16A1 | 9.82E-36 | 1.36E-34 |
| TNFAIP6 | 5.03E-35 | 6.91E-34 |
| GPC3 | 8.89E-35 | 1.21E-33 |
| CNTNAP2 | 7.04E-34 | 9.51E-33 |
| RCN3 | 1.47E-33 | 1.97E-32 |
| REEP6 | 1.89E-33 | 2.50E-32 |
| LTBP2 | 6.00E-32 | 7.89E-31 |
| MGST1 | 1.05E-31 | 1.37E-30 |
| EMILIN1 | 4.46E-31 | 5.77E-30 |
| PRRX1 | 4.72E-30 | 6.05E-29 |
| C1S | 2.28E-29 | 2.90E-28 |
| ANGPTL2 | 4.06E-25 | 5.12E-24 |
| TTC36 | 2.69E-23 | 3.36E-22 |
| HGD | 4.08E-23 | 5.06E-22 |
| DACT1 | 4.89E-23 | 6.02E-22 |
| FBLN2 | 4.28E-22 | 5.22E-21 |
| NID2 | 4.84E-22 | 5.87E-21 |
| APOD | 4.18E-20 | 5.03E-19 |
| TIMP1 | 6.96E-20 | 8.31E-19 |
| CD79A | 9.22E-20 | 1.09E-18 |
| EFEMP1 | 4.39E-19 | 5.16E-18 |
| MMP13 | 5.02E-19 | 5.85E-18 |
| PDGFRL | 2.50E-18 | 2.90E-17 |
| CKMT1A | 4.14E-18 | 4.76E-17 |
| CILP | 6.08E-18 | 6.93E-17 |
| GRP | 7.05E-18 | 7.98E-17 |
| P4HA3 | 9.17E-18 | 1.03E-16 |
| TTC6 | 4.05E-17 | 4.52E-16 |
| PSCA | 6.80E-17 | 7.53E-16 |
| MS4A1 | 9.98E-17 | 1.10E-15 |

| | | |
|---|---|---|
| PODNL1 | 1.82E-16 | 1.99E-15 |
| CRISPLD2 | 2.35E-16 | 2.55E-15 |
| GLT8D2 | 1.11E-15 | 1.20E-14 |
| PDPN | 4.11E-15 | 4.39E-14 |
| CD79B | 1.35E-14 | 1.44E-13 |
| WNT7B | 1.84E-13 | 1.94E-12 |
| COL14A1 | 2.01E-13 | 2.10E-12 |
| TNFRSF13C | 2.84E-13 | 2.96E-12 |
| LPAR1 | 6.11E-13 | 6.31E-12 |
| PODN | 1.76E-12 | 1.81E-11 |
| MSX2 | 2.79E-12 | 2.84E-11 |
| SPIB | 3.32E-12 | 3.36E-11 |
| MFAP4 | 9.99E-12 | 1.00E-10 |
| CCDC160 | 1.58E-11 | 1.58E-10 |
| DHRS2 | 2.08E-11 | 2.06E-10 |
| S100A7 | 7.66E-11 | 7.55E-10 |
| RHBDL1 | 7.78E-11 | 7.63E-10 |
| NOX4 | 9.17E-11 | 8.93E-10 |
| IGFBP6 | 1.61E-10 | 1.56E-09 |
| CD19 | 3.19E-10 | 3.07E-09 |
| SPTSSB | 1.20E-09 | 1.15E-08 |
| ALDH1A1 | 1.21E-09 | 1.15E-08 |
| ZMYND10 | 1.33E-09 | 1.26E-08 |
| IGF1 | 2.54E-09 | 2.38E-08 |
| GXYLT2 | 3.02E-09 | 2.82E-08 |
| IGHM | 5.76E-09 | 5.34E-08 |
| ANK2 | 1.16E-08 | 1.07E-07 |
| FBLN5 | 1.25E-08 | 1.15E-07 |
| ADAM33 | 1.36E-08 | 1.24E-07 |
| SCG5 | 1.45E-08 | 1.31E-07 |
| CXCR5 | 2.04E-08 | 1.84E-07 |
| VPREB3 | 7.76E-08 | 6.91E-07 |
| HLA.DOB | 7.77E-08 | 6.91E-07 |
| LAMA2 | 1.03E-07 | 9.15E-07 |
| FCRL5 | 1.16E-07 | 1.02E-06 |
| WISP2 | 4.49E-07 | 3.93E-06 |
| MCF2L.AS1 | 1.06E-06 | 9.26E-06 |
| FCRLA | 1.17E-06 | 1.01E-05 |
| DPT | 1.37E-06 | 1.17E-05 |
| ADAM28 | 2.20E-06 | 1.88E-05 |
| SRPX | 2.38E-06 | 2.02E-05 |
| HVCN1 | 2.52E-06 | 2.13E-05 |
| FCRL2 | 3.59E-06 | 3.02E-05 |
| BANK1 | 4.92E-06 | 4.12E-05 |
| TINCR | 5.10E-06 | 4.24E-05 |
| STRA6 | 1.08E-05 | 8.96E-05 |
| ECM2 | 1.35E-05 | 1.11E-04 |
| LY6D | 1.36E-05 | 1.11E-04 |

| | | |
|---|---|---|
| FCRL1 | 5.31E-05 | 4.32E-04 |
| DNM3OS | 8.22E-05 | 6.66E-04 |
| HRASLS | 1.00E-04 | 8.09E-04 |
| PNOC | 1.02E-04 | 8.18E-04 |
| CLGN | 1.08E-04 | 8.59E-04 |
| GAPT | 1.09E-04 | 8.62E-04 |
| NR2F1 | 1.16E-04 | 9.16E-04 |
| FST | 1.49E-04 | 1.17E-03 |
| CCDC8 | 2.41E-04 | 1.88E-03 |
| RUNX1T1 | 2.54E-04 | 1.97E-03 |
| LRRC17 | 3.25E-04 | 2.51E-03 |
| CCL11 | 4.09E-04 | 3.14E-03 |
| ABCA6 | 8.04E-04 | 6.16E-03 |
| ABCA8 | 1.34E-03 | 1.02E-02 |
| SCG2 | 2.58E-03 | 1.96E-02 |
| LINC00926 | 3.22E-03 | 2.43E-02 |
| IGHD | 3.30E-03 | 2.48E-02 |
| ZNF750 | 3.57E-03 | 2.67E-02 |
| TNFRSF13B | 3.66E-03 | 2.72E-02 |
| SLIT2 | 5.27E-03 | 3.90E-02 |
| ABCA9 | 6.00E-03 | 4.42E-02 |
| BMPR1B | 7.16E-03 | 5.25E-02 |
| OSR1 | 8.94E-03 | 6.49E-02 |
| CSMD2 | 8.94E-03 | 6.49E-02 |
| PLA2G10 | 1.09E-02 | 7.88E-02 |
| VGLL3 | 1.29E-02 | 9.30E-02 |
| CNTN4 | 1.31E-02 | 9.37E-02 |
| KLK1 | 1.46E-02 | 1.04E-01 |
| PAPLN | 1.52E-02 | 1.08E-01 |
| CLCA2 | 1.55E-02 | 1.09E-01 |
| PLEKHH2 | 1.84E-02 | 1.29E-01 |
| LSAMP | 1.85E-02 | 1.29E-01 |
| FAM129C | 2.00E-02 | 1.39E-01 |
| ABCA10 | 1.99E-02 | 1.39E-01 |
| MUCL1 | 2.43E-02 | 1.68E-01 |
| HAS2 | 2.48E-02 | 1.70E-01 |
| STAP1 | 2.58E-02 | 1.77E-01 |
| RGMA | 3.02E-02 | 2.05E-01 |
| LPAR4 | 3.55E-02 | 2.41E-01 |
| CAPN6 | 6.75E-02 | 4.56E-01 |
| ADH1C | 8.51E-02 | 5.72E-01 |
| TBX18 | 9.64E-02 | 6.45E-01 |
| XIRP1 | 9.76E-02 | 6.51E-01 |
| ENPP6 | 1.18E-01 | 7.80E-01 |
| KHDRBS2 | 9.12E-01 | 1.00E+00 |
| AC022182.1 | 2.32E-01 | 1.00E+00 |
| ABCB4 | 6.91E-01 | 1.00E+00 |
| KLHL14 | 2.69E-01 | 1.00E+00 |

| | | |
|---|---|---|
| MEIS2 | 4.04E-01 | 1.00E+00 |
| TCEAL7 | 9.41E-01 | 1.00E+00 |
| MMP16 | 2.80E-01 | 1.00E+00 |
| MAB21L1 | 1.86E-01 | 1.00E+00 |
| IGDCC4 | 1.92E-01 | 1.00E+00 |
| PLA2G5 | 2.01E-01 | 1.00E+00 |
| TCEAL5 | 6.83E-01 | 1.00E+00 |
| XPNPEP2 | 1.66E-01 | 1.00E+00 |
| TFAP2B | 3.76E-01 | 1.00E+00 |
| LINC01133 | 7.00E-01 | 1.00E+00 |
| FABP7 | 2.03E-01 | 1.00E+00 |
| NCCRP1 | 3.25E-01 | 1.00E+00 |
| LINC01285 | 5.29E-01 | 1.00E+00 |
| DKK1 | 1.54E-01 | 1.00E+00 |
| MSLN | 3.85E-01 | 1.00E+00 |
| LEMD1 | 2.49E-01 | 1.00E+00 |

### Table S5: Detailed information of datasets

| | |
|---|---|
| **Human breast cancer** | |
| Data source (scRNA) | https://www.ncbi.nlm.nih.gov/geo/query/acc.cgi?acc=GSE176078 |
| Data source (Spatial) | https://www.10xgenomics.com/datasets/human-breast-cancer-block-a-section-1-1-standard-1-0-0 |
| **Human heart** | |
| Data source (scRNA) | https://support.10xgenomics.com/spatial-gene-expression/datasets/1.1.0/V1_Human_Heart |
| **Human pancreas** | |
| Data source (scRNA) | https://www.ncbi.nlm.nih.gov/geo/query/acc.cgi?acc=GSE84133 |
| **Lung Adenocarcinoma** | |
| Data source (scRNA) | https://www.ncbi.nlm.nih.gov/geo/query/acc.cgi?acc=GSE131907 |
| **Mouse olfactory** | |
| Data source (scRNA) | https://www.ncbi.nlm.nih.gov/geo/query/acc.cgi?acc=GSE148360 |
| Data source (Spatial) | https://github.com/CaiGroup/seqFISH-PLUS |
| **PBMC** | |
| Data source (scRNA) | https://support.10xgenomics.com/single-cell-gene-expression/datasets/1.1.0/pbmc3k |

### Simulation Study

To validate the effectiveness of the correlated gating mechanism in selecting informative genes while accounting for feature correlation, we conducted a controlled simulation study.

We generated a synthetic dataset with $n$ =1000 samples and $p$ =10 informative features. Labels were generated based only on the first two features, while the remaining features were noise. To introduce a correlation structure, we simulated nine additional features for each of the original ten features, each highly correlated with its corresponding original feature. This expanded the dataset to a total of 100 features.

After applying the correlated binary gating mechanism, we observed that scGPD accurately prioritized the true causal features, while the correlated features are constrained to be selected. This demonstrates that the binary gating mechanism effectively discourages the selection of redundant information, promoting a compact and non-redundant gene panel.

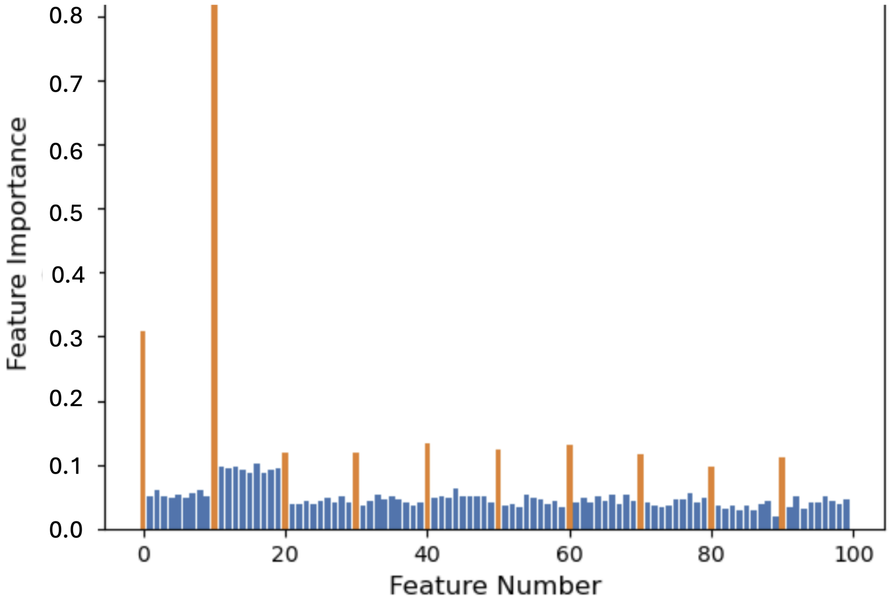


Figure S4: Simulation results.
